# Bacterial defense via RES-mediated NAD^+^ depletion is countered by phage phosphatases

**DOI:** 10.64898/2026.01.28.702374

**Authors:** Ilya Osterman, Bohdana Hurieva, Erez Yirmiya, Dina Hochhauser, Maxim Itkin, Sergey Malitsky, Rotem Sorek

## Abstract

Many bacterial defense systems restrict phage infection by breaking the molecule NAD^+^ to its constituents, adenosine diphosphate ribose (ADPR) and nicotinamide (Nam). To counter NAD^+^ depletion-mediated defense, phages evolved NAD^+^ reconstitution pathway 1 (NARP1), which uses ADPR and Nam to rebuild NAD^+^. Here we report a bacterial defense system called aRES, involving RES-domain proteins that degrade NAD^+^ into Nam and ADPR-1’’-phosphate (ADPR-1P). This molecule cannot serve as a substrate for NARP1, so that NAD^+^ depletion by aRES defends against phages even if they encode NARP1. We further discover that some phages evolved an extended NARP1 pathway capable of overcoming aRES defense. In these phages, the NARP1 operon also includes a specialized phosphatase, which dephosphorylates ADPR-1P to form ADPR, a substrate from which NARP1 then reconstitutes NAD^+^. Other phages encode inhibitors that directly bind aRES proteins and physically block their active sites. Our study describes new layers in the NAD^+^-centric arms race between bacteria and phages and highlights the centrality of the NAD^+^ pool in cellular battles between viruses and their hosts.

## Introduction

Nicotinamide adenine dinucleotide (NAD^+^) is a molecule essential for cellular energy metabolism. Studies from recent years show that this molecule plays a central role in bacterial defense against phage infection^1,2^. Many bacterial defense systems deplete NAD^+^ from the cell once phage infection is sensed, depriving the phage of this essential molecule, and, sometimes, leading to premature cell lysis^3,4^. NAD^+^ depletion during phage infection can lead to the death or growth arrest of the infected cell, but since the infecting phage cannot replicate in the absence of NAD^+^, phage propagation is prevented^3^.

Multiple types of defense systems were shown to degrade NAD^+^ in response to phage infection. A partial list includes CBASS^5^, Pycsar^6^, type I Thoeris^4^, prokaryotic Argonautes^3,7^, DSR1 and DSR2^3^, bacterial NLR-like Avast proteins^8^, the Nezha^9^ and Kongming^10^ systems, and more. In all these systems, the enzymatic domains SIR2 (SIRim), TIR, or SEFIR break NAD^+^ by cleaving the N-glycosidic bond between the nicotinamide ring and the ribose, producing the products ADP-ribose (ADPR) and free nicotinamide^3,6,11^. It was estimated that more than 15% of all bacterial genomes encode at least one defense system whose immune function involves NAD^+^ depletion^1^.

To counteract NAD^+^ depletion, phages have evolved specific enzymatic pathways that rebuild NAD^+^. The most abundant of these pathways is NAD^+^ Reconstitution Pathway 1 (NARP1)^12^. This two-enzyme pathway rebuilds NAD^+^ from ADPR and nicotinamide via a two-step process: First, the enzyme ADPR-pyrophosphate synthetase (Adps) converts ADPR to ADPR-pyrophosphate (ADPR-PP) using ATP as the pyrophosphate donor (Figure S1). Then, the enzyme nicotinamide ADPR transferase (Namat) replaces the pyrophosphate with nicotinamide to reconstitute NAD^+12^. A second NAD^+^ reconstitution pathway in phages, NARP2, uses canonical NAD^+^ salvage enzymes to rebuild NAD^+^ from phosphoribosyl pyrophosphate (PRPP), ATP, and nicotinamide^12,13^. As NARP1 uses ADPR and nicotinamide, which are the direct products of NAD^+^ cleavage by the SIR2, TIR and SEFIR domains, it does not build NAD^+^ in infected cells unless an NAD^+^-depleting system has been activated, and hence NARP1 does not inflict surplus metabolic burden for the phage^12^. NARP2, on the other hand, constructs excess NAD^+^ in infected cells regardless of whether they employ NAD^+^ depletion, thus adding a metabolic load during phage infection^12^ (Figure S1). Possibly, the advantages of NARP1 caused it to be more common in nature: this pathway is encoded on at least 4% of phages as compared to 1% of phages encoding NARP2 ^12^.

In this study, we describe a new defense system that relies on a RES domain, which cleaves NAD^+^ between the nicotinamide ring and the ADPR ribose, but leaves a phosphate group on ADPR to generate ADPR-1’’-phosphate (ADPR-1P)^14,15^. We show that this mode of NAD^+^ cleavage renders phage NARP1 inactive, because the first enzyme in the pathway (Adps) cannot use ADPR-1P as a substrate. We further demonstrate that certain phages evolved to encode a dedicated phosphatase in the NARP1 operon, which dephosphorylates ADPR-1P to yield ADPR, thereby rescuing NAD^+^ synthesis via NARP1. This reveals a new layer of the arms race dynamics between bacterial defense and phage counter-defense, expanding our understanding of both NAD^+^ metabolism and phage-host interactions.

## Results

### aRES proteins protect bacteria from phage via NAD^+^ depletion

We initiated this study by focusing on a family of proteins that are frequently present in defense islands in bacterial genomes (Figure 1A). These proteins caught our attention because they encoded a RES domain, which was previously shown to possess NAD^+^-degrading activity in toxin-antitoxin systems implicated in bacterial persistence^14-17^. The RES-domain proteins we studied were not operonically associated with other proteins, suggesting that they function as standalone proteins and not within a toxin-antitoxin system (Figure 1A). We cloned three members of this family, one from *Escherichia coli* UC5, one from *Enterobacter chengduensis*, and a third from *Klebsiella pneumoniae*, and expressed each of them in *E. coli* MG1655. All three were able to protect bacteria against infection by diverse phages (Figure 1B), and protection was associated with bacterial growth arrest (Figure 1C). We named this new family of defensive proteins aRES, for <u>a</u>ntiphage <u>RES</u>-domain proteins. Ares is also the name of one of the twelve Olympian gods in Greek mythology, so designating the new system as aRES follows the convention of naming antiphage systems after mythological deities^11,18^.

**Figure 1.**
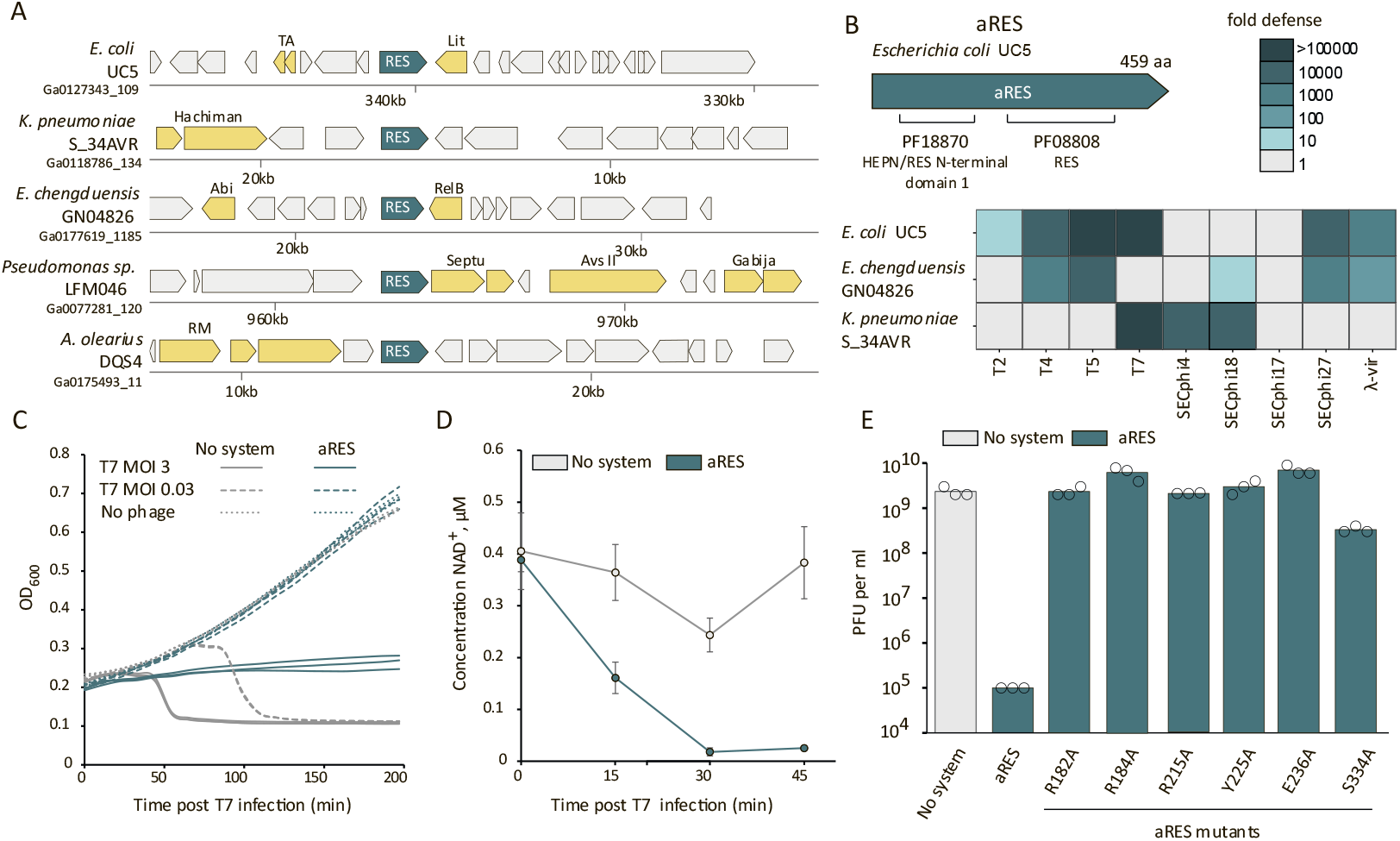
aRES protects bacteria from phages via NAD^+^ depletion. **A.** A stand-alone RES-domain protein is found in defense islands. Representative instances of the system are presented in their genomic environments. Genes known to be involved in defense are shown in yellow. RM, restriction modification; TA, toxin-antitoxin, Abi, abortive infection. The encoding strain and the respective DNA scaffold accession in the IMG database^20^ are indicated to the left. **B**. aRES protects against phages. Schematic representation of the aRES gene from *Escherichia coli* UC5, comprising two protein domains. aRES from *Escherichia coli* UC5, *Klebsiella pneumoniae* S_34AVR and *Enterobacter chengduensis* GN04826 strains were expressed in *E. coli* MG1655. Fold defense was quantified by serial dilution plaque assays, comparing the efficiency of plating phages on the system-containing strain to the efficiency of plating on a control strain that lacks the systems and contains an empty vector instead. Data represent an average of three replicates. PFU quantification for individual replicates is presented in Figure S3. **C**. Growth curves of *E. coli* MG1655 cells expressing the *E. coli* UC5 aRES or an empty vector, infected by phage T7 at a multiplicity of infection(MOI) of 0.03 or 3 (or 0 for uninfected cells). Data from three biological replicates are presented as individual curves. **D**. NAD^+^ concentration in lysates obtained from cells expressing the *E. coli* UC5 aRES, or control cells with an empty vector (no system). Samples were analyzed 0, 15, 30 and 45 minutes after infection with T7 at an MOI 3. Average of 3 replicates, error bars represent standard deviation. **E**. Mutations in aRES abolish defense against phage. Data represent PFUs per milliliter of T7 phage infecting cells that express WT *E. coli* UC5 aRES or aRES mutated in the indicated residues. Bar graphs are the average of three independent replicates, with individual data points overlaid.

Given that some bacterial toxins with RES domains are known to exert their toxicity via NAD^+^ depletion^16,19^, we hypothesized that aRES defense is also associated with depletion of NAD^+^. Indeed, quantification of intracellular NAD^+^ levels showed that NAD^+^ is depleted in aRES-containing cells during infection (Figure 1D). Point mutations in residues that were predicted, based on a structural alignment with RES toxins, to form the RES active sites, abolished defense, suggesting that the catalytic function of the RES domain is essential for phage resistance (Figures 1E, S2).

### NARP1 fails to overcomes aRES defense

Previous studies on RES domain toxins involved in persistence of bacterial pathogens showed that the RES enzyme cleaves NAD^+^ at the nicotinamide–ribose N-glycosidic bond, with inorganic phosphate acting as the nucleophile^14,15^. The inorganic phosphate becomes attached to the ADPR product, so that the products of the NAD^+^ cleavage reaction are nicotinamide and ADPR-1P^14,15^. Our data show that the levels of ADPR-1P sharply rise in aRES-expressing cells during phage infection, but this molecule is not detected in infected cells not expressing aRES (Figure 2A). In agreement with these results, incubation of NAD^+^ with total lysates from phage-infected, aRES-expressing cells, resulted in complete NAD^+^ degradation and the appearance of ADPR-1P as the major product (Figure 2B). These results suggest that the RES domain in the aRES defense system functions as an NAD^+^ phosphorylase, similar to its activity within toxin-antitoxin systems^14,15^ (Figure 2C).

**Figure 2.**
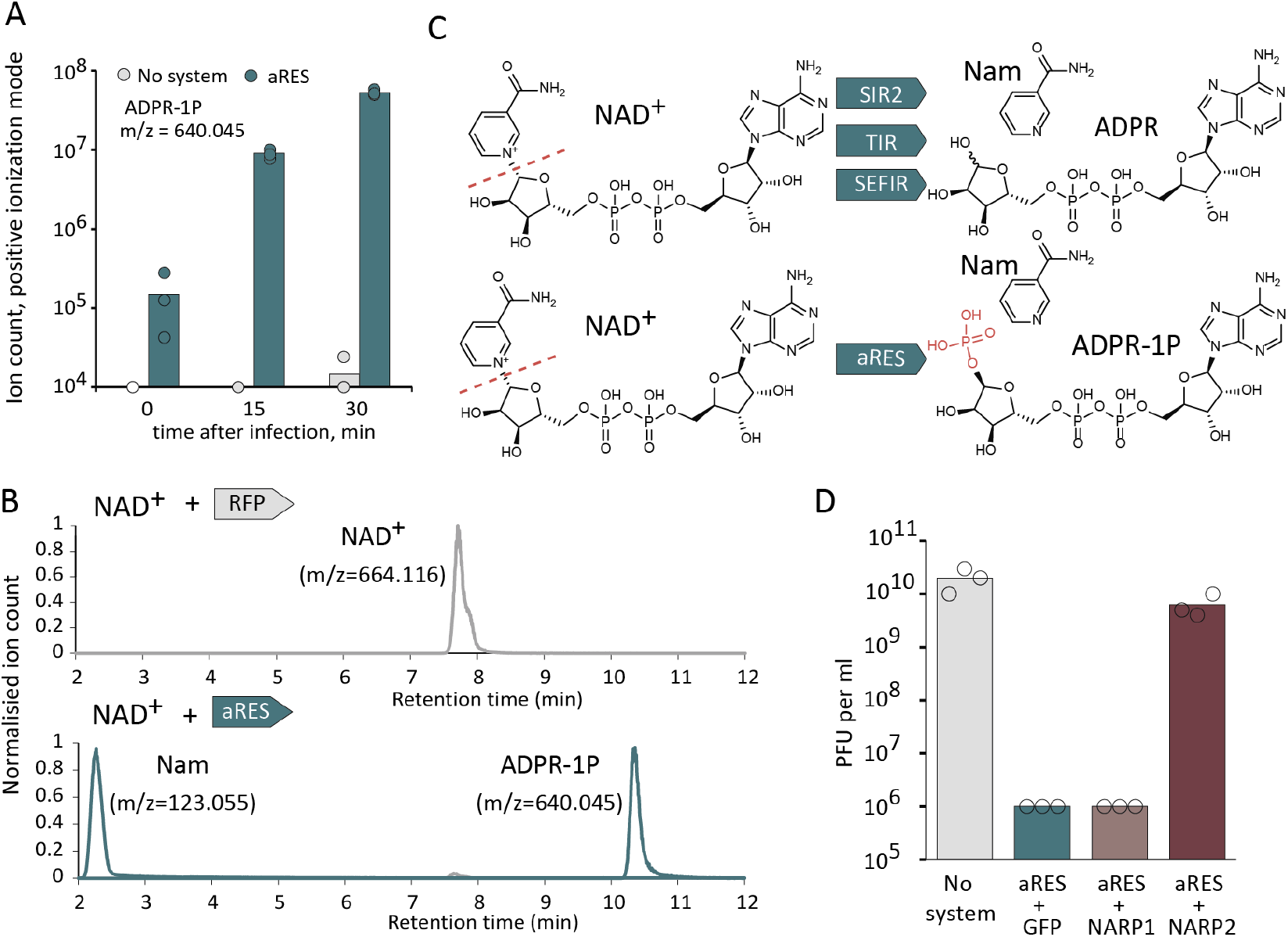
NARP1 does not overturn aRES defense. **A.** ADPR-1P accumulates in cells expressing aRES during infection with phage T7. Cells were infected at MOI = 3, measurements taken at 15 or 30 min post infection, or prior to infection (time 0). Presented are untargeted LC-MS-derived ion count data; bars represent the mean area under the curve of three experiments, with individual data points overlaid. **B**. LC-MS analysis of the product of NAD^+^ degradation by a cell lysate derived from aRES-expressing cells that were infected with phage T7. Produced molecules were identified by comparing the peaks with the retention times of chemical standard molecules (Figure S4). **C**. Schemes of NAD^+^ cleavage by SIR2, TIR, SEFIR and aRES defense proteins. **D**. Anti-defense effect of NARP1, NARP2 or GFP co-expressed in *E. coli* together with aRES from *E. coli* UC5. Bacteria were infected by phage T7, and 10-fold serial dilution plaque assays were performed. Bar graphs are the average of three independent replicates, with individual data points overlaid.

Given that the most common phage NAD^+^ reconstitution pathway, NARP1, requires ADPR as a substrate, we hypothesized that NARP1 will not be able to rescue phages from the consequences of aRES defense. Consistent with this hypothesis, NARP1 did not subdue the ability of aRES to defend against phage, while NARP2 efficiently canceled aRES defense (Figure 2D). These results show that aRES is an NAD^+^-depleting defense system that can overcome the NARP1 counter-defense measures of phages. The results may also explain why encoding NARP2, which relies on PRPP rather than on ADPR as a substrate, is preferred over encoding NARP1 in some phages, despite the additional metabolic burden it inflicts on the phage (Figure S1).

### A phage phosphatase restores NARP1 NAD^+^ reconstitution activity

While examining NARP1 operons in phage genomes, we noticed that in some cases these operons contain additional genes beyond the two core NARP1 genes. One of the most common genes associated with NARP1 was a gene encoding a protein with a Macro domain, a domain known to interact with ADPR derivatives^21,22^ (Figure 3A; Table S1). We found that 15% of phages carrying NARP1 also included this protein associated with the NARP1 operon. Homology searches via HHpred^23^ showed that the closest homolog with a known function is the Poa1p protein from the yeast *Saccharomyces cerevisiae* (Figure S5). Poa1p is involved in yeast tRNA splicing, and is known to be a phosphatase that converts ADPR-1P into ADPR^24^. We reasoned that supplementation of NARP1 with an enzyme that dephosphorylates ADPR-1P to ADPR will render the extended NARP1 pathway active against aRES-mediated NAD^+^ depletion.

**Figure 3.**
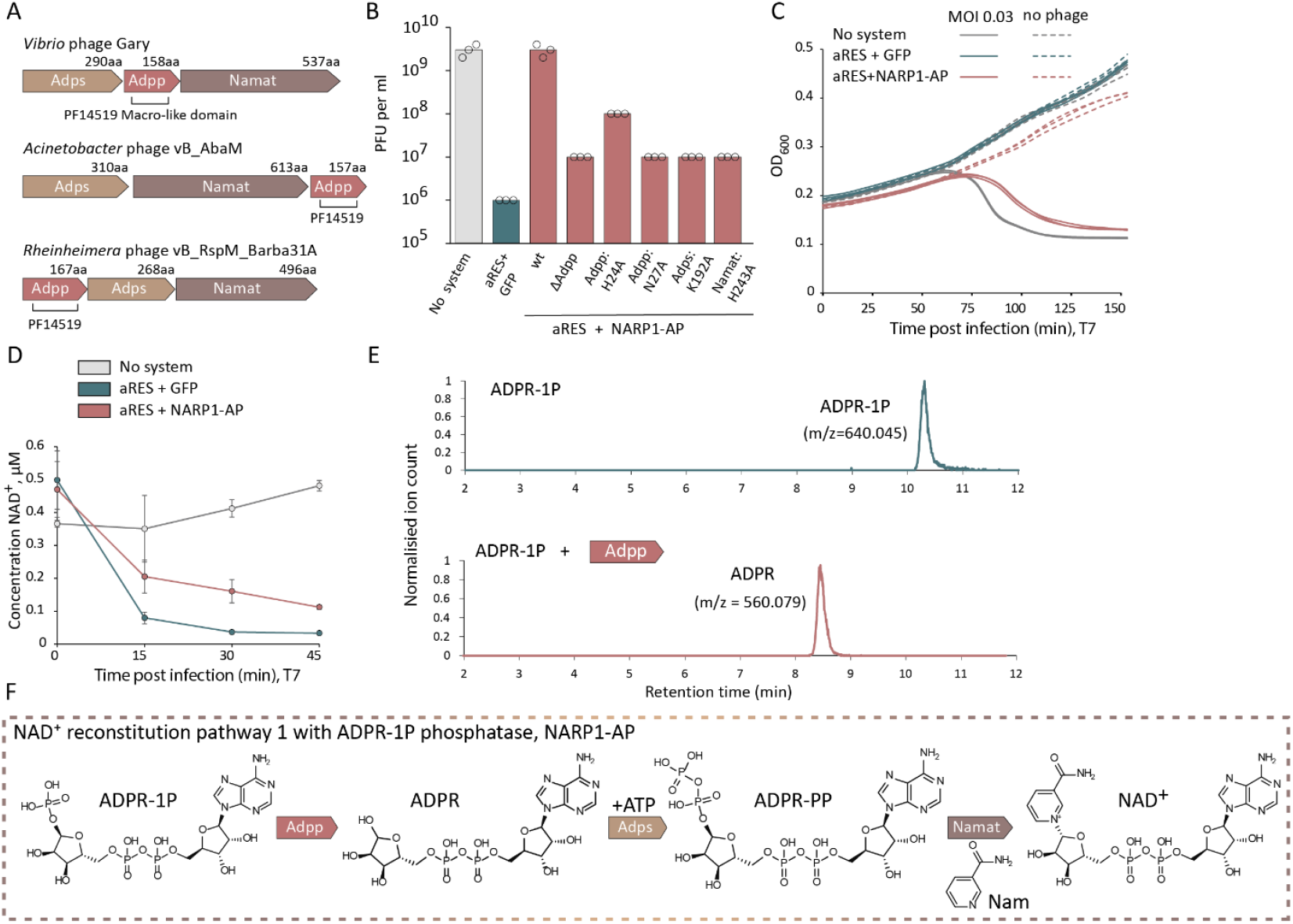
A NARP1-associated phosphatase converts ADPR-1P to ADPR. **A.** Schematic representation of NARP1 operons with an associated Macro-domain protein (Adpp). **B**. Mutations in the Marco-domain protein Adpp abolish anti-defense against aRES. Data represent PFUs per milliliter of phage T7 infecting cells that express aRES with WT or mutated variants of NARP1-AP. Bar graphs are the average of three independent replicates, with individual data points overlaid. **C**. Growth curves of *E. coli* MG1655 cells expressing aRES with GFP or NARP1-AP, or an empty vector, infected by phage T7 at an MOI of 0.03 or 0 for uninfected cells. Data from three biological replicates are presented as individual curves. **D**. NAD^+^ concentration in lysates obtained from cells expressing aRES with GFP or NARP1-P, or control cells with an empty vector instead (no system). Samples were analyzed 0,15,30 and 45 minutes after infection with T7 at an MOI 3. Average of 3 replicates, error bars represent standard deviation. **E**. A purified Macro-domain Adpp protein from *Vibrio phage* Gary was incubated with ADPR-1P. Shown are LC-MS ion count data. Representative of 3 replicates. ADPR was identified by comparing the peaks with the retention time of the chemical standard (Figure S4). **F**. Schematic representation of the NARP1-AP pathway.

To test this hypothesis, we cloned the 3-gene operon Adps-phosphatase-Namat from *Vibrio* phage Gary, and co-expressed it with aRES. We observed that the operon was able to override aRES defense (Figures 3B, 3C). Inactivation of any of the three genes, or deletion of the predicted phosphatase, abolished the anti-defense phenotype, indicating that all enzymatic components are required (Figure 3B). NAD^+^ measurements in cell lysates confirmed that expression of the 3-gene operon resulted in elevated NAD^+^ levels in infected cells despite the presence of aRES (Figure 3D).

To examine if the phage homolog of Poa1p is indeed an ADPR-1P phosphatase, we purified this protein from the NARP1 operon of *Vibrio* phage Gary and incubated it with ADPR-1P. LC–MS analysis verified that this enzyme converted ADPR-1P to ADPR (Figure 3E). Together, these results show that the extended NARP1 pathway, which includes Adps, Namat and ADPR-1P phosphatase (now called Adpp to conform with NARP enzymes nomenclature), overcomes aRES defense. We name this pathway NARP1 with an associated phosphatase, or NARP1-AP for short (Figure 3F). Notably, NARP1-AP retains all functional features of NARP1 and can counteract not only aRES but also other defense systems that convert NAD^+^ into ADPR and Nam. For example, NARP1-AP was also able to cancel defense by the Nezha system (Figure S6).

### A family of phage anti-defense proteins that bind and inhibit aRES

We found homologs of the aRES defensive protein in about 4% of bacterial and archaeal genomes (1,526 genomes out of 38,149 analyzed; Figure S7, Tables S2, S3). Given the abundance of aRES in microbial genomes, we hypothesized that phages might have evolved additional strategies to overcome aRES-mediated defense. A recent study demonstrated that large-scale application of Alphafold-Multimer enables the discovery of phage proteins that inhibit bacterial defense proteins via physical binding^25^. In this strategy, a defense protein of choice is serially co-folded with proteins from a phage protein database, and high-scoring hits are considered reliable if homologs of the phage protein also co-fold with the defense protein with high scores^25^. To search for aRES inhibitors within phage proteomes, we applied this strategy using a set of ∼133,000 short phage proteins of unknown function that represent a clustered database of ∼112 million phage proteins from IMG/VR v4^26^. This allowed us to identify a family of phage proteins whose members were consistently predicted to bind aRES with high Alphafold3 confidence scores, and in which binding involved a flexible loop that interacts with the aRES active site pocket (Figure 4A, B, C). To test if proteins from this family can inhibit aRES defense in vivo, we co-expressed seven such proteins together with the aRES protein. Six of the tested proteins completely abolished aRES defense (Figure S7 B,C). We name this protein family aRad1 (aRES anti-defense 1).

**Figure 4.**
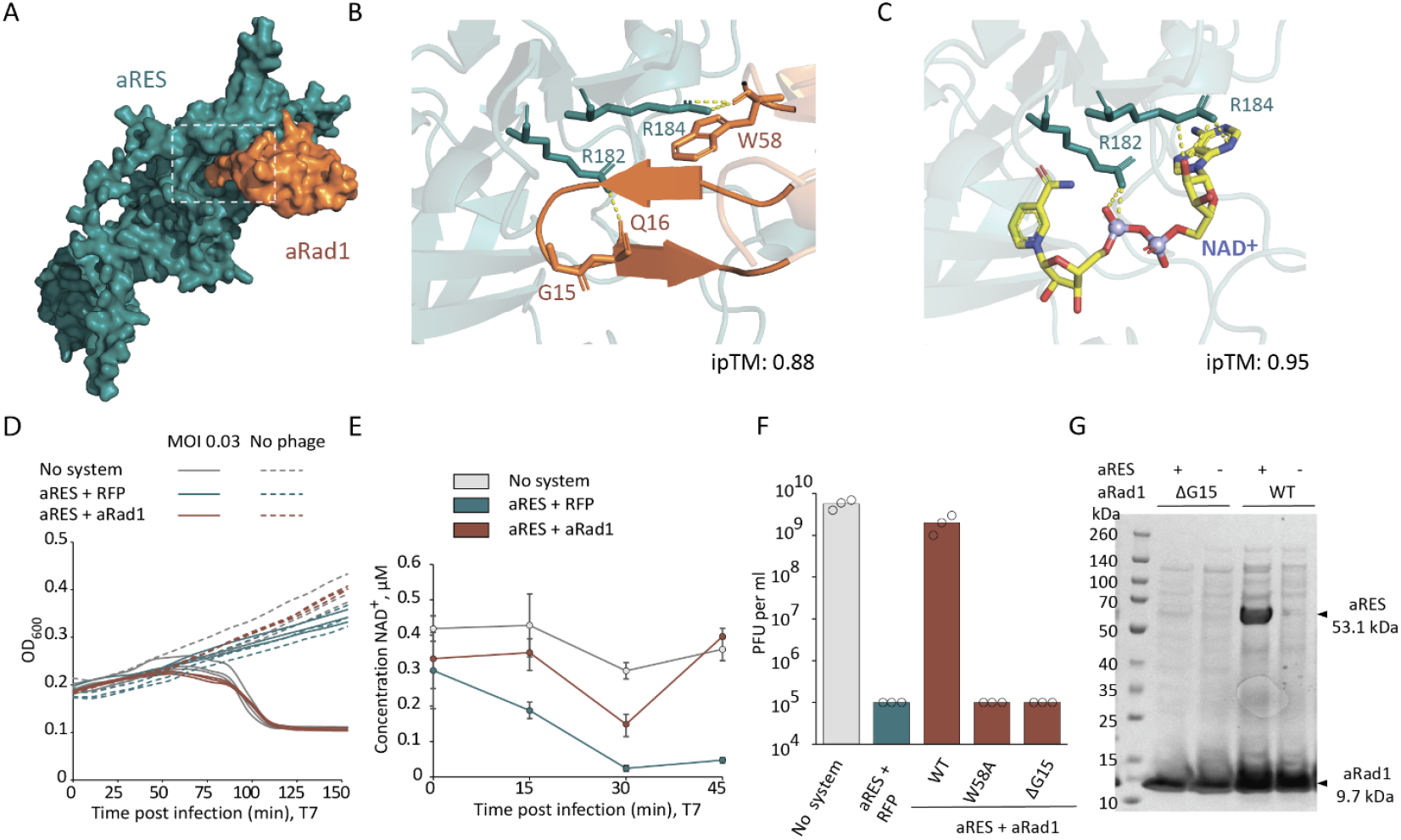
aRad1 binds and inhibits aRES. **A.** AlphaFold3-predicted structure of aRES from *E. coli* UC5 in complex with aRad1 (aRad1b). The active site region of aRES is marked with a dashed box. **B**. Close-up view of the Alphafold3-predicted interactions between the aRad1 loop and the aRES active site. **C**. Close-up view of the Alphafold3-predicted NAD^+^ binding site in aRES. Yellow dashed lines indicate predicted hydrogen bond interactions. **D**. Growth curves of *E. coli* MG1655 cells co-expressing the *E. coli* UC5 aRES with RFP or with aRad1, or an empty vector, infected by phage T7 at an MOI of 0.03 or 0 for uninfected cells. Data from three biological replicates are presented as individual curves. **E**. NAD^+^ concentration in lysates obtained from cells expressing aRES with RFP or aRad1, or control cells with an empty vector instead (no system). Samples were analyzed 0,15,30 and 45 minutes after infection with T7 at an MOI 3. Average of 3 replicates, error bars represent standard deviation. **F**. Mutations in aRad1 abolish anti-defense against aRES. Data represent PFUs per milliliter of phage T7 infecting cells that express aRES with WT or mutated variants of aRad1 in the indicated residues. Bar graphs are the average of three independent replicates, with individual data points overlaid. **G**. Pulldown of a His-tagged aRad1 (WT or the ΔG15 mutant) following co-expression with aRES. WT aRad1, but not the mutated version, co-elutes with aRES. Shown is an SDS-PAGE of proteins following pulldown. Representative of 3 replicates.

aRad1 is a small protein with no predicted function annotation. In liquid cultures of cells expressing aRES, phage infection was successful if aRad1 was also expressed. This phenotype was accompanied by inhibition of NAD^+^ degradation, supporting that aRad1 inhibits the enzymatic activity of aRES (Figure 4D, E). AlphaFold3 predicts aRad1 to bind the RES domain of aRES via an extended interface, which includes a flexible loop that penetrates the active site of the RES domain (Figure 4A). The flexible loop, and a conserved tryptophan residue from another section of the protein, overlaps the NAD^+^ binding pocket of the RES domain (Figure 4B, C). A point mutation in the conserved tryptophan residue, or deletion of a conserved glycine residue in the flexible loop, abolished the anti-defense activity of aRad1 (Figure 4F; Figure S7).

To test whether aRad1 inhibition of aRES involves direct protein-protein interaction, we co-expressed aRES with a tagged version of aRad1 and used affinity chromatography to pull down aRad1. We observed that aRad1 co-purified with aRES, indicating complex formation between the two proteins (Figure 4G). The aRad1 variant mutated in the flexible loop pulled down aRES only weakly, indicating that this loop participates in the physical interaction between the proteins. Together, these results show that aRES defense can be neutralized by a small phage-encoded protein that binds to the active site of the RES domain and blocks its NAD^+^-depletion activity.

## Discussion

Studies from recent years have exposed a central role for NAD^+^ manipulation in bacterial immune systems^1-8^. While most NAD^+^-depleting systems in bacteria rely on enzymes that cleave the nicotinamide ring to generate Nam and ADPR, our discovery of aRES identifies an alternative mode for depleting NAD^+^ in the context of bacterial defense (Figure 5). Furthermore, in a recent study^27^ we demonstrated a third mechanism for immune-related NAD^+^ depletion, in which calcineurin-domain proteins in the Metis defense system cleave NAD^+^ between the two phosphate moieties, generating the molecules AMP and NMN (Figure 5). It was recently estimated that 15.9% of all bacteria carry one of the “canonical” NAD^+^-depleting immune pathways^1^, but our discoveries of aRES and Metis now suggest that the fraction of bacteria defending against phages via NAD^+^ depletion exceeds 20%.

**Figure 5.**
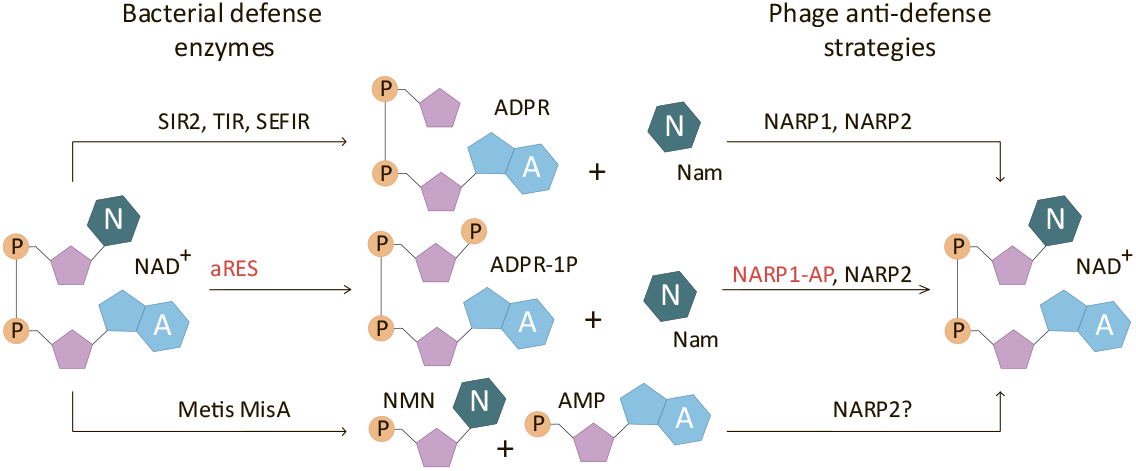
NAD^+^ depletion and reconstitution in bacterial defense and phage counter defense.

The existence of defense systems that cleave NAD^+^ in a non-canonical manner can explain the diversity of NAD^+^ reconstitution pathways in phages^12,13,28^. When countering canonical NAD^+^ depletion, NARP1 is more efficient than NARP2 because it rebuilds NAD^+^ from the direct products of NAD^+^ cleavage, and thus does not produce excess NAD^+^ unless NAD^+^ depletion has taken place^12^. However, the reliance on ADPR as substrate is also the Achilles heel of NARP1, as our data show it cannot rescue the phage if the defense system cleaves NAD^+^ in a non-canonical manner.

Our results show that phages developed two different metabolic strategies to overcome non-canonical NAD^+^ depletion. First, they can encode NARP2, a pathway that synthesizes excess NAD^+^ regardless of whether it has been depleted, and is hence metabolically wasteful and somewhat toxic^12^. Second, they can enhance NARP1 by adding a dedicated phosphatase that removes the phosphate placed by aRES on ADPR following NAD^+^ cleavage. This method retains NARP1 efficiency, but is specific to countering aRES defense and will likely not work against defense systems that cleave NAD^+^ differently (Figure 5).

In addition to metabolic reconstitution of NAD^+^ via NARP1-AP, we also showed that phages can encode aRad1 proteins that bind the active site of aRES and block its NADase activity. The mode of action of aRad1 is similar to that of the Thoeris anti-defense proteins Tad3, Tad4, Tad5, Tad6 and Tad7, all of which inhibit bacterial immunity by binding the defense protein and blocking its NAD^+^-processing active site via a flexible loop^25^.

The phosphatase we discovered in the NARP1-AP pathway has a Macro domain, an ADPR-handling domain common in cellular organisms ^21,22^. Notably, Macro domain proteins are also widespread among eukaryotic viruses^29^, including coronaviruses^30^. It was previously demonstrated that Macro domain proteins from viruses infecting humans (hepatitis E virus and severe acute respiratory syndrome coronavirus) can remove the phosphate group from ADPR-1P *in vitro*^31^, raising the intriguing possibility that viruses use these proteins to counter host defense, and that animal antiviral pathways may manipulate NAD^+^ in a manner similar to bacterial immune systems.

Until a few years ago, the role of NAD^+^ in bacterial immunity and phage counter-immunity was almost completely unknown. Our study joins a flurry of recent studies that demonstrate the importance of NAD^+^ manipulation in phage-host interactions^1^. NAD^+^ is not considered central to animal immunity, although a few observations suggest a role for NAD^+^ depletion in animals. For example, an NAD^+^-processing TIR-domain protein in *C. elegans* was implicated in antimicrobial immunity^32^, and the a TIR-STING protein in the mollusk *C. gigas* was suggested to deplete NAD^+^ in response to cGAS-like signaling^5^. If future studies show that NAD^+^ depletion is common in animal antiviral immunity akin to bacteria, it would not be surprising if animal viruses also encode strategies to counter such defense.

## Acknowledgements

We thank members of the Sorek lab for constructive discussions during this study. R.S. was supported, in part, by the European Research Council (grant ERC-AdG GA 101018520), the Israel Science Foundation (MAPATS grant 2720/22), the Deutsche Forschungsgemeinschaft (SPP 2330, grant 464312965), the Minerva Foundation with funding from the Federal German Ministry for Education and Research, and a research grant from Magnus Konow in honor of his mother Olga Konow Rappaport. I.O. was supported by the Ministry of Absorption’s New Immigrant program. E.Y. was supported by the Clore Scholars Program and, in part, by the Israeli Council for Higher Education (CHE) via the Weizmann Data Science Research Center.

## Supplementary Figures

**Supplementary Figure S1.**
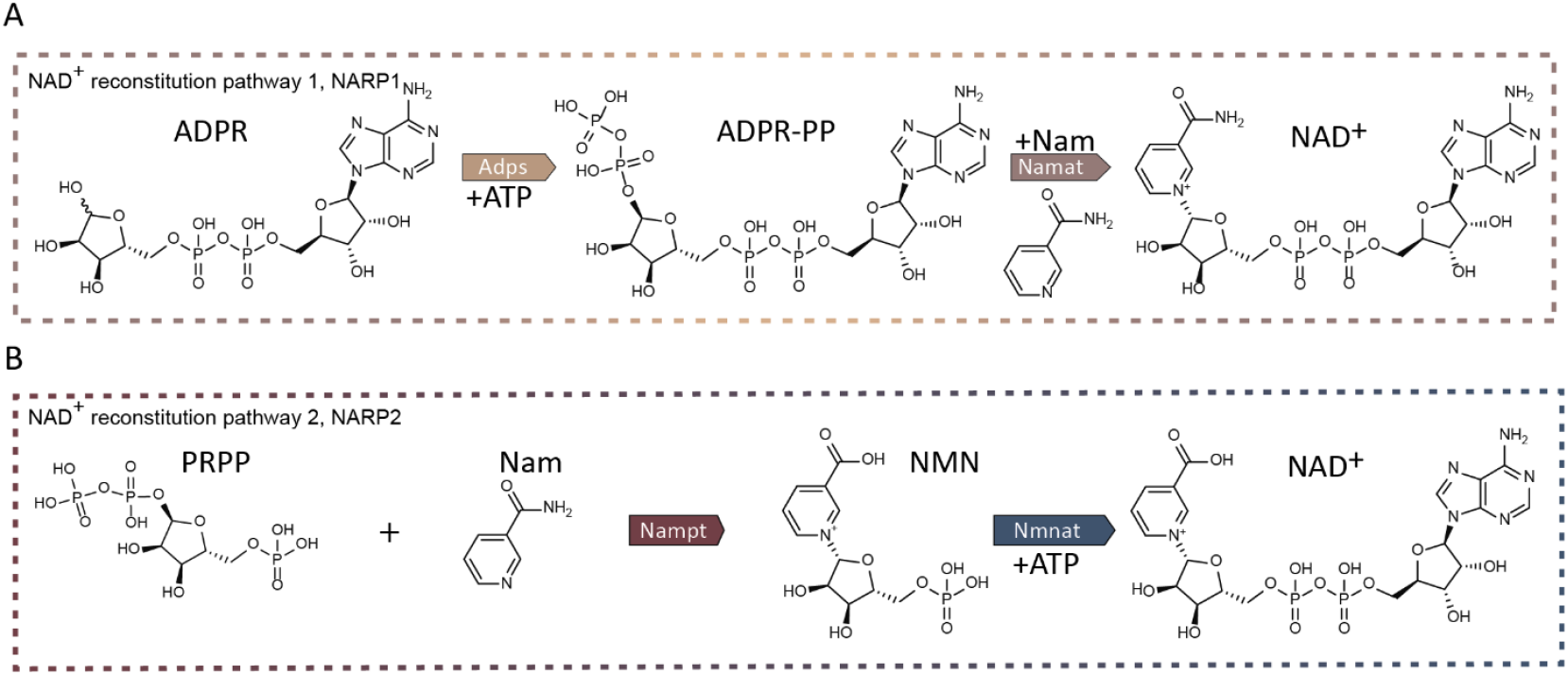
Schematic representation of the NARP1 (A) and NARP2 (B) NAD^+^ reconstitution pathways.

**Supplementary Figure S2.**
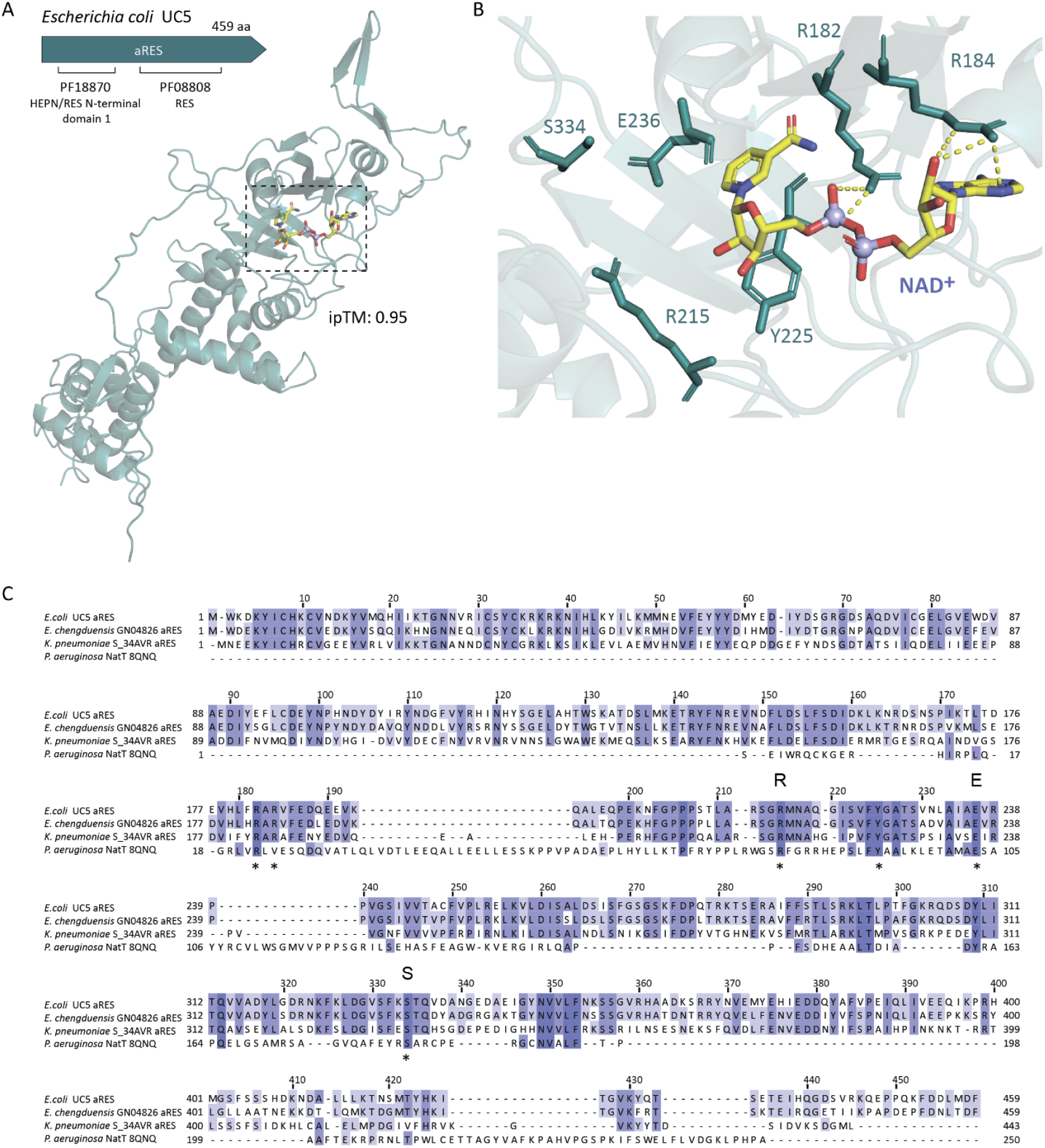
The aRES protein. **A.** AlphaFold3-predicted structure of aRES from *E. coli* UC5 in complex with NAD^+^. The active site region of aRES is marked with a dashed box. **B**. Close-up view of the predicted NAD^+^ binding site. Residues mutated in this study are highlighted in dark green. Yellow dashed lines indicate predicted hydrogen bond interactions. **C**. Structure-guided sequence alignment^33^ of aRES proteins together with the RES toxin NatT^14^. The indicative R, E, and S residues are marked by capital letters. NAD^+^-interacting amino acids mutated in this study are marked by asterisks.

**Supplementary Figure S3.**
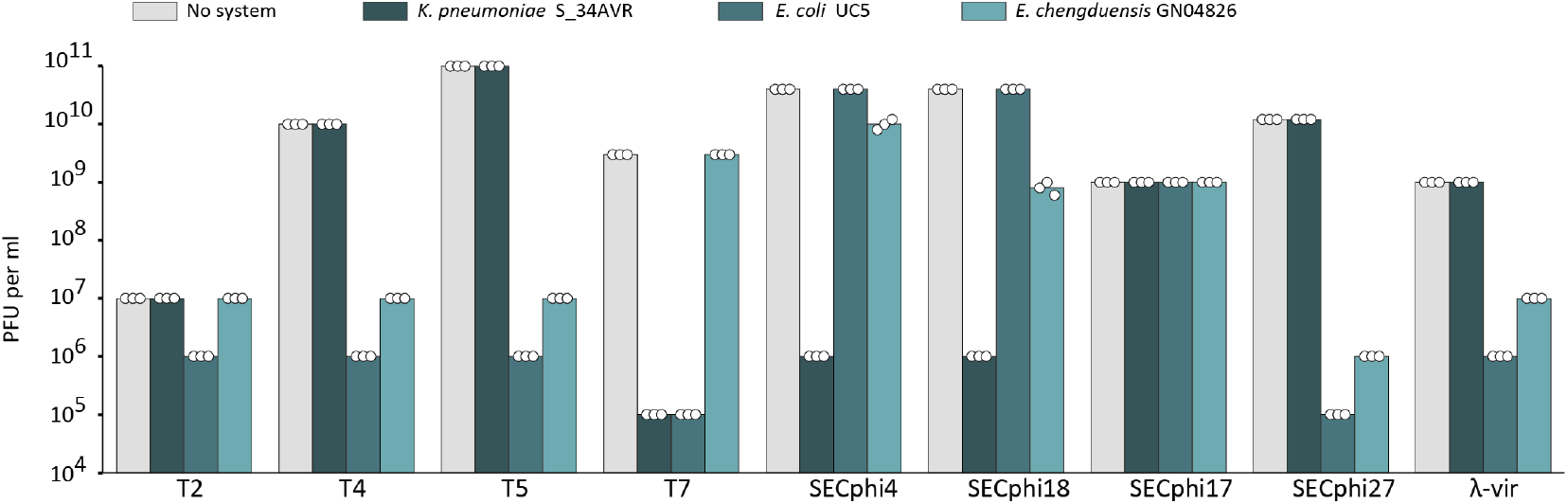
Serial dilution plaque assay experiments. Data represent PFUs per milliliter of phages infecting *E. coli* MG1655 control cells (no system) and cells expressing aRES systems cloned from *Klebsiella pneumoniae* S_34AVR, *Escherichia coli* UC5, and *Enterobacter chengduensis* GN04826. Bar graphs represent the average of three independent replicates, with individual data points overlaid.

**Supplementary Figure S4.**
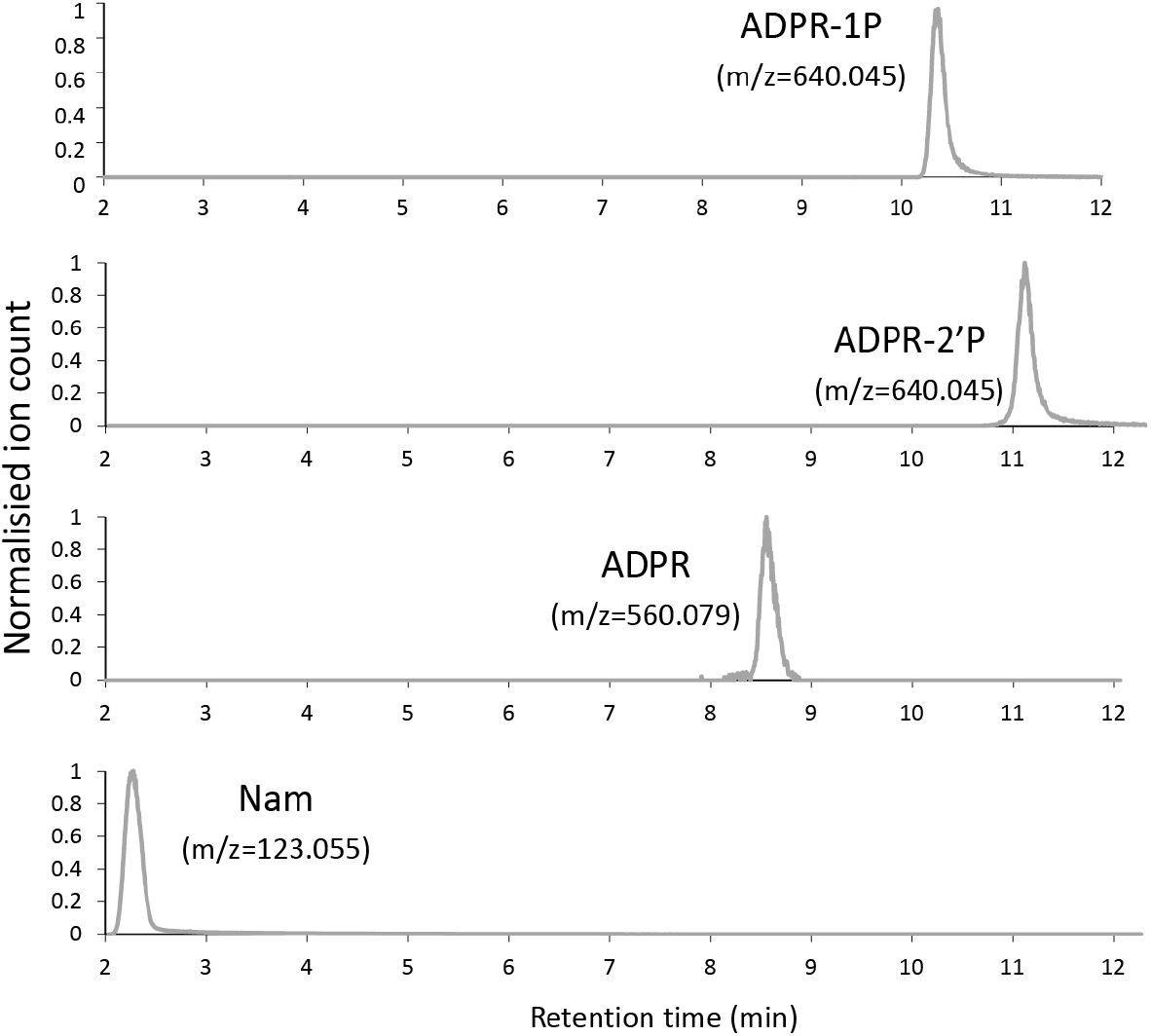
LC-MS analysis of chemical standards of ADPR-2P, ADPR and Nam. ADPR-1P was purified as indicated in the Methods section. Data for ADPR-1P also appear in Figure 2B.

**Supplementary Figure S5.**
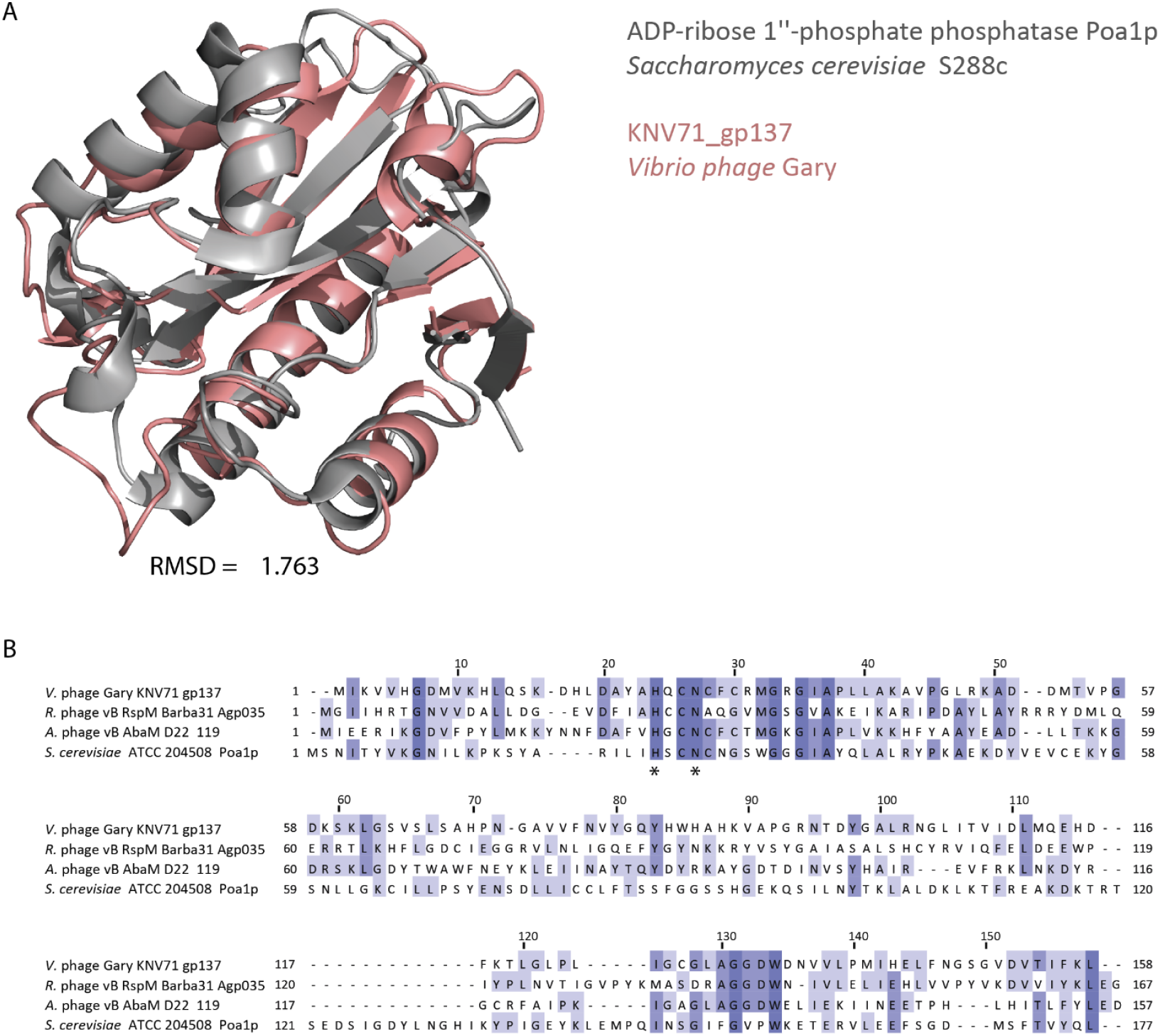
The NARP1-associated Macro-domain protein is homologous to yeast Poa1p. A. AlphaFold3-predicted structures of protein KNV71_gp137 from *Vibrio phage* Gary aligned with *Saccharomyces cerevisiae* Poa1p (6LFQ)^34^. **B**. Sequence alignment of NARP1-associated Macro-domain proteins from phages and Poa1p from *S. cerevisiae*. Amino acids mutated in this study are marked with asterisks.

**Supplementary Figure S6.**
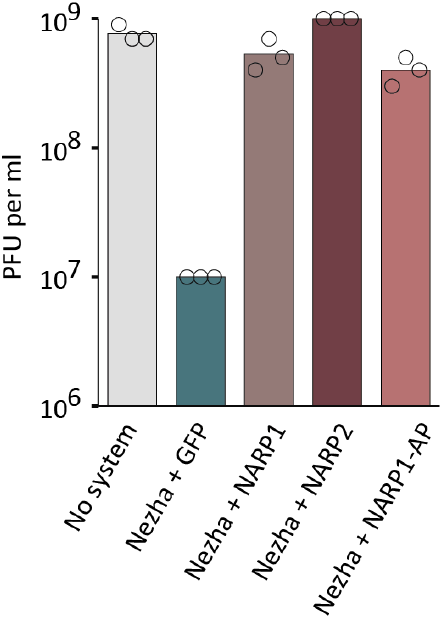
Anti-defense effect of NARP1, NARP2 and NARP1-AP. NARP1 from *Bacillus* phage SPβL1, NARP2 from phage *Vibrio* Phage KVP40, and NARP1-AP from phage *Vibrio* phage Gary were co-expressed in *E. coli* together with the Nezha defense system from *Paenibacillus sp*. 453MF^3^. Data represent PFUs per milliliter of phage vB_EcoM-KAW1E185 on the indicated strains. Bar graphs are the average of three independent replicates, with individual data points overlaid.

**Supplementary Figure S7.**
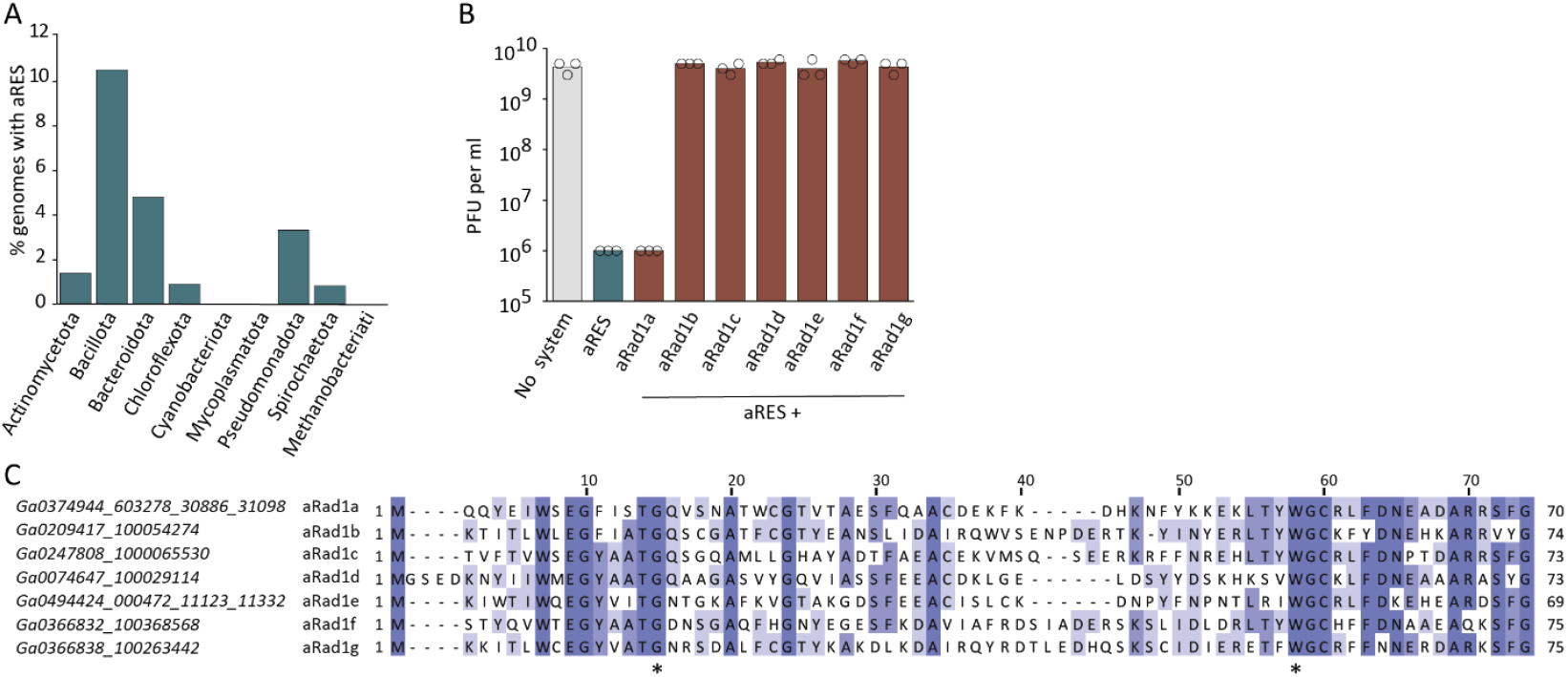
The aRES defense and aRAD anti-defense protein families. **A.** Fraction of genomes encoding aRES genes across bacterial and archaeal phyla, shown for phyla with at least 200 genomes in the analyzed dataset. **B**. Anti-defense effect of proteins from the aRad1 family or GFP (control) co-expressed in *E. coli* together with *E. coli* UC5 aRES. Bar graphs are the average of three independent replicates, with individual data points overlaid. **C**. Sequence alignment of aRad1 homologs. Residues mutated in this study are marked by asterisks. Accession numbers of aRad1 homologs in the IMG/VR database^26^ are indicated.

## Methods

### Strains and growth conditions

All *E. coli* strains were grown in MMB media (lysogeny broth (LB) supplemented with 0.1 mM MnCl2 and 5 mM MgCl2) at 37°C with 200 rpm shaking or on solid 1.5% LB agar plates. Ampicillin 100 μg/mL, chloramphenicol 30 μg/mL, streptomycin 50 μg/mL or kanamycin 50 μg/mL were added when necessary for plasmid maintenance. *E. coli* DH5a (NEB) was used for cloning, BL21 (DE3) for protein purification and MG1655 for experiments with phages. All chemicals were obtained from Sigma Aldrich unless stated otherwise. All phages used in the study were amplified from a single plaque at 37°C in *E. coli* MG1655 culture in MMB until the culture collapsed. A list of all plasmids, strains and phages used in this study can be found in Supplementary Table S4.

### Plasmid construction and transformation

DNA amplification for cloning was performed using KAPA HiFi HotStart ReadyMix (Roche) according to the manufacturer’s instructions. All primers were obtained from Sigma Aldrich. Supplementary Table S4 lists all primers used in this study.

For cloning of large fragments, PCR products with 20-nucleotide overlaps were generated and treated with FastDigest DpnI restriction enzyme (ThermoFisher) for 30 min at 37°C. Fragments were then Gibson-assembled by NEBuilder HiFi DNA Assembly Master Mix (NEB) according to the manufacturer’s instructions and used for transformation into DH5a (NEB #C2987H). Single colonies were checked by PCR and plasmids were validated by a plasmid sequencing service (Plasmidsaurus). Plasmids ordered from Twist Bioscience or GenScript Corporation are listed in Supplementary Table S4. Verified plasmids were used for the transformation of *E. coli* strains using the standard TSS protocol^35^. aRES from *Escherichia coli* UC5 was integrated into the genome of *E. coli* MG1655 using the Tn7 integration plasmid (downstream of the gmlS gene, spectinomycin resistance, induced by anhydrotetracycline, Sigma cat.37919)^36^. Genome integration was verified by a genome sequencing service (Plasmidsaures).

### Plaque assays

Phage titer was determined as described previously^37^. 300 µL of the overnight bacterial cultures were mixed with 30 mL of melted MMB 0.5% agar, poured on 10 cm square plates and left to dry for 1 h at room temperature. L-Arabinose was added to a concentration of 0.2% to induce gene expression from pBAD plasmids. IPTG was added to a concentration of 1 mM to induce expression from pBba6c plasmid. Tenfold dilutions of phages were prepared in MMB and 10 µL of each dilution was dropped onto the plates. Plates were incubated overnight at 25°C. Plaque-forming units were counted the next day.

### Liquid infection assay

Overnight bacterial cultures of *E. coli* MG1655 were diluted in MMB (1:100) and grown until reaching an optical density at 600nm (OD_600_) of 0.3 at 25°C. Then, 180 µL of cultures were transferred to a 96-well plate and infected with 20 µL of phages at various MOIs. Culture growth was followed by OD_600_ measurements every 10 min on a Tecan Infinite 200 plate reader at 25°C. Strains, plasmids and inducer concentrations are indicated in Supplementary Table S4.

### Cell lysates preparation

*E. coli* MG1655 cells with integrated aRES gene from *E. coli* UC5 system or a negative control lacking the system were grown at 25 °C, 200 rpm until reaching an OD_600_ of 0.3 in 150 mL of MMB. Cells were then infected with phage T7 with MOI=3. Samples were collected before infection, 15, 30 and 45 min after infection. At each time point, 50 mL of cells were centrifuged for 10 min at 4°C, 4000 *g*, and the pellet was frozen in liquid nitrogen and stored at -80°C. To extract cell metabolites from frozen pellets, the pellet was resuspended in 600 μL of 60% ice-cold ethanol. Samples were transferred to FastPrep Lysing Matrix B in a 2 mL tube (MP Biomedicals, cat #116911100) and lysed at 4°C using a FastPrep bead beater for two rounds of 40 s at 6 ms^−1^. The tubes were then centrifuged at 4°C for 10 min at 15,000 *g*. The supernatant was then transferred to an Amicon Ultra-0.5 Centrifugal Filter Unit 3 kDa (Merck Millipore, no. UFC500396) and centrifuged for 45 min at 4°C, 12,000 *g*. Lysates were analyzed by LC-MS.

### LC-MS analysis of the lysates

Metabolic profiling of polar metabolites within filtered cell lysates was carried out as described previously^38^ with minor modifications as described below. In brief, analysis was carried out using an Acquity I class UPLC System combined with a mass spectrometer Q Exactive Plus Orbitrap (Thermo Fisher Scientific), which was operated in positive ionization mode. The LC separation was carried out using the SeQuant Zic-pHilic (150 mm × 2.1 mm) with the SeQuant guard column (20 mm × 2.1 mm; Merck). Mobile phase B was acetonitrile and mobile phase A was 20 mM ammonium carbonate with 0.1% ammonia hydroxide in deionized distilled water/acetonitrile (80:20, v/v). The flow rate was kept at 200 μL/min, and the gradient was as follows: 0–2 min 75% of B, 14 min 25% of B, 18 min 25% of B, 19 min 75% of B, for 4 min, 23 min 75% of B. Peak areas were extracted using MZmine 2 with an accepted deviation of 5 ppm. All standards used were obtained from Sigma.

### NAD^+^ detection assay

*E. coli* MG1655 cells carrying a defense system or negative control cells lacking the system were grown at 25 °C, 200 rpm until reaching an OD_600_ of 0.3. Strains, plasmids and inducer concentrations are indicated in Supplementary Table S4. Then, 180 µL of cultures were transferred to a 96-well plate and infected with 20 µL of phages at an MOI of 3. Plates were incubated at 25°C with shaking in a TECAN Infinite200 plate reader. 15 µL of cells were taken before phage infection (t=0) and at 15, 30 and 45 min post-infection. Cells were mixed with 20 µL of 100% ethanol and kept at -20°C. For the NAD^+^ detection assay, samples were diluted 1:5 in 100 mM sodium phosphate buffer (pH 7.5). Then 5 µL of the samples were mixed with 5 µL of NAD/NADH-Glo™ detection reagent (Promega corp.) in a Nunc^®^ 384-well polystyrene plate (Sigma p5991-1CS). Plates were incubated at 25°C in a TECAN Infinite200 plate reader and luminescence was measured every 10 min.

### Protein purification

Adpp from *Vibrio* phage Gary was cloned into the pET28-His-bdSUMO expression vector (TWIST Bioscience, USA), which encodes an N-terminal His14-bdSUMO tag^39^. Plasmids were transformed into LOBSTR-BL21(DE3)-RIL cells (Kerafast, USA), and cells were grown in 50 mL of MMB media at 37°C until mid-log phase (OD_600_∼0.5), then IPTG was added to a concentration of 0.5 mM and cells were further grown overnight at 16°C, centrifuged and flash-frozen. His-tagged proteins were purified using 50 µL NEBExpress® Ni-NTA Magnetic Beads (NEB) according to the manufacturer’s protocol. Elution was done using 1 µg His-tagged bdSENP1 protease in 100 µL of cleavage buffer (20 mM HEPES 125 KCl and 1 mM DTT), for 12 hours at 4°C with intensive shaking.

### NAD^+^ degradation by a cell lysate derived from aRES-expressing cells that were infected with phage T7

*E. coli* MG1655 cells with integrated aRES gene from *E. coli* UC5 and negative control harboring RFP gene were grown at 25 °C, 200 rpm until reaching an OD_600_ of 0.3 in 50 mL of MMB. Cells were then infected with phage T7 with MOI=3. 30 min after infection, 50 mL of cells were centrifuged for 10 min at 4°C, 4000 *g* for 10 min, and the pellet was frozen in liquid nitrogen and stored at -80°C. To prepare a cell lysate, the pellet was resuspended in 600 μL of 100 mM sodium phosphate buffer (pH 7.5). Samples were transferred to FastPrep Lysing Matrix B in a 2 mL tube (MP Biomedicals, cat #116911100) and lysed at 4°C using a FastPrep bead beater for two rounds of 40 s at 6 ms^−1^. The tubes were then centrifuged at 4°C for 10 min at 15,000 *g*. The supernatant was then transferred to an Amicon Ultra-0.5 Centrifugal Filter Unit 3 kDa (Merck Millipore, no. UFC500396) and centrifuged for 45 min at 4°C, 12,000 *g*. Concentrated lysate was diluted 1:10 in 100 mM sodium phosphate buffer (pH 7.5) and again filtered on Amicon Ultra-0.5 Centrifugal Filter Unit 3 kDa. This procedure was repeated 3 times to wash off cellular metabolites. Finally, the lysate was diluted to 600 µL of 100 mM sodium phosphate buffer, and 100 µL of activated lysate was mixed with 1mM of NAD^+^. After incubation for 2 hours at room temperature, the products of the reaction were filtered on an Amicon Ultra-0.5 Centrifugal Filter Unit 3 kDa and analyzed by LC-MS with the same conditions as the lysates.

### Purification of ADPR-1P

Products of the reaction from the previous step were separated by HPLC using Phenomenex Luna 5 µm C18(2) 100 Å HPLC Column. The following protocol was used for all runs: 2 min of mobile phase A 100%, 1 min 75% A and 25% B, 1 min 50% A and 50% B, 1 min 20% A and 80% B, 6 min 100 % A and 4 min 100% A, 1 mL/min flow rate. Mobile phase A was 20 mM potassium phosphate pH 6 and B was 20 mM potassium phosphate pH 6 in 20% methanol. Peak of ADPR-1P was collected and verified by LC-MS. Concentration of ADPR-1P was estimated by normalization to the absorbance (A_260_) of ADPR solution with known concentration (100 μM).

### Enzymatic activity of Adpp

100 µM of ADPR-1P was incubated with 1 µM of the purified MisB in 10 mM Tris-HCl buffer (pH=8) with supplementation of 10 mM of MgCl_2_ for 120 min at 25°C. The products of the reactions were diluted 1:4 in water and were transferred to an Amicon Ultra-0.5 Centrifugal Filter Unit 3 kDa (Merck Millipore, no. UFC500396), centrifuged for 45 min at 4°C, 12,000 *g*, and analyzed by LC-MS with the same parameters as cell lysates.

### Structure prediction

AlphaFold3^40^ with default parameters was used to generate structural models for aRES, Adpp and aRad1 together with the respective ligand. The following SMILES code was used for NAD^+^:

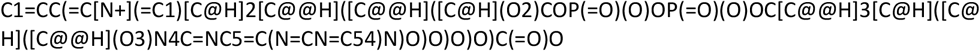

### Detection of Macro domain proteins in NARP1 operons

A database of ∼1.8 million genes derived from 20,185 phage genomes was downloaded from the INPHARED database^41^ on 1 May 2023. NARP1 pathways, defined by the presence of Adps and Namat genes in close proximity, were identified as previously described^12^, resulting in 859 NARP1 pathways. All protein-coding genes located within two genes upstream or downstream of each NARP1 system were scanned with HHpred^42^ against the Pfam database^43^. Proteins annotated with Pfams PF14519, PF01661, or PF20016 were considered as Macro-domain containing.

### Detection of putative aRES systems

A database of ∼135 million genes derived from ∼38,000 bacterial and archaeal genomes was downloaded from the Integrated Microbial Genomes (IMG)^44^database in October 2017. All protein sequences were aggregated into protein clusters as previously described^11^.

Proteins were considered candidate aRES proteins if they were annotated in IMG with the Pfam^43^ annotation PF00808, or if they belonged to protein clusters in which PF00808 was the most abundant Pfam annotation among cluster members. Candidates were further filtered to include only proteins with lengths between 300 and 600 amino acids. Each remaining protein sequence was analyzed using the hhblits option of hhsuite (version 3.2.0)^42^ to generate multiple sequence alignments (MSAs) using the UniRef30 database and with the parameters “-oa3m -n 3 -M first”. The resulting MSAs were then searched with the hhsearch option of hhsuite against the Pfam database (version 35) using the parameters “-p 80 - loc -z 1 -b 1 -ssm 2 -sc 1 -seq 1”. Only sequences with hits to both Pfam domains PF00808 and PF18870 with probabilities above 80% were retained as final aRES genes.

### aRad prediction using AlphaFold3-Multimer

5.5 million phage scaffolds labeled as “high-confidence virus” were downloaded from the IMG/VR v4 database^26^. The corresponding protein sequences were clustered using the “cluster” option of MMseqs2^45^ with default parameters. For each cluster containing at least 30 non-identical members, a representative sequence was extracted, and its structure was predicted using AlphaFold2^46^ with default parameters, yielding 182,179 phage protein structures. Clusters representing proteins with an average size above 300 amino acids, or proteins with an HHPred^23^ hits to an annotated protein of known function, were discarded.

AlphaFold3^40^ was then used with default parameters to co-fold each remaining phage sequence with the RES domain extracted from IMG^44^ accession 2753914621. The 10 phage proteins with the highest ipTM scores when co-folded with the RES domain were examined manually. One phage sequence (IMG ID Ga0374944_603278_30886_31098), which displayed a flexible loop entering the RES active-site pocket, was selected for downstream analysis, as similar structural features were previously shown to be predictive of phage-encoded anti-defense proteins^25^.

The predicted structure of Ga0374944_603278_30886_31098 was next searched against the set of 182,179 phage protein structures using Foldseek release 5.53465f065^47^ with default parameters. Hits with a probability score above 90% were collected together with all members of their respective MMseqs2 clusters, resulting in a set of 156 homologs.

Each of the 156 homologs was then co-folded with each RES domain from the following IMG accessions: 2508547945, 2574243703, 2703310322, 2729545185, and 2753914621. For every RES-phage protein pair, five AlphaFold-Multimer predictions were generated using the five different AF3 models, and the interaction score was defined as the average ipTM across the five models. Seven phage homologs that exhibited high scores when co-folded with multiple RES domains were selected for experimental validation.

